# In-vivo egfp expression in the honeybee *Apis mellifera* induced by electroporation and viral expression vector

**DOI:** 10.1101/2022.01.31.478532

**Authors:** Gérard Leboulle, Nora Gehne, Anja Froese, Randolf Menzel

## Abstract

In this study we describe egfp expression induced by two techniques: in vivo electroporation and viral transduction in several cell types of the adult honeybee brain. Non-neuronal and neuronal cell types were identified and the expression persisted at least during three days. Kenyon cells, optic lobe neurons and protocerebral lobe neurons were electroporated. Astrocyte-like glia cells, fibrous lamellar glia cells and cortex glia cells were identified. Viral transduction targeted one specific type of glia cells that could not be identified. EGFP positive cells were rather variable after electroporation, and viral transduction resulted in more homogenous groups of positive cells. We propose that these techniques remain a good alternative to transgenic animals because they potentially target only somatic cells.

## INTRODUCTION

The honeybee is a model in neurobiology providing access to the study of neural processes underlying perception, learning and memory in the context of remarkable behavioural capabilities [1, 2] [3]. However, until recently efficient tools to selectively manipulate its brain function for behavioural studies were mostly lacking. The genome revolution decrypted its genome and opened new perspectives in socio-genomics by relating gene expression profiles to social organisation and physiological states [4] [5] [6]

The development of the antisense oligo and RNA interference techniques allowed a break-through by solving partially the specificity issues associated with pharmacological treatment, but they were not always associated with robust effects (i.e. small amplitude and short duration of the interference on gene expression inhibition), which prevents them from becoming a routine techniques for honeybee neurobiological studies [7] [8, 9]. These limitations can be explained since the injected molecules did not penetrate efficiently into the cells. Disposing of tools to more efficiently introduce genetic material into the cell should help to overcome these problems. Moreover, these techniques allowed only to downregulate gene expression.

Early attempts to generate transgenic animals by adding recombinant DNA to drone sperm before the inoculation of a queen were presented [10]. Unfortunately, this study did not allow a breakthrough in the generation of transgenic animals. Only recently transgenic animals were generated by injecting eggs with the transposable element Piggybac and egfp (enhanced green fluorescent protein) expression could be induced in the whole organism [11]. More recently the CRISPR/Cas9 technique was successfully applied to knock down the expression of several genes [12] [13] [14]. These techniques have the disadvantage that germinal cell lines are modified, which hinder the release of these animals in their natural environment for ethical reasons. The animals have to be confined into the laboratory or in flying cages, which strongly limits the experimental range.

In this regard, techniques modifying only somatic cells should allow circumventing these problems. In vivo electroporation relies on the generation of field strength to modify the organisation of the cell membrane and induce the formation of pores allowing exogenous DNA to penetrate into the cell [15]. The first successful attempts of in vivo electroporation in honeybees reported egfp expression after injection of plasmidic DNA controlling gene expression by a pCMV promoter [16]. The vector used was non-integrative, which further limited the biological risk because the expression of the exogene was transient. The technique was used later to evaluate in vivo the functionality of vectors engineered with native honeybee promoters [17]. In the context of in vivo electroporation, cell type specific promoters do not only allow a precise control of the physiological manipulation, they also increase the safety of the system by limiting the cell types expressing the exogene. Moreover, neurons are postmitotic cells and although neurogenesis has been demonstrated in several species [18], it is not reported in honeybee workers [19], and it probably does not lead to the generation of new egg cells in queens, thus further limiting the biological risk.

Viral expression vectors were also widely used in neuroscience to modify neurons [20]. They rely on the fusion of viral particles with the cell membrane to allow passing the genetic material into the cell.

Although previous works on electroporation demonstrated efficient expression of egfp, they did not describe precisely the electroporated cell types [16, 17]. In this study, we compared different electroporation protocols for their potential to induce egfp expression in the honeybee brain under the control of the pCMV (cytomegalovirus promoter) promoter. We showed that the expression increased during the days following the electroporation of the plasmid. We also showed that glia cells were predominantly electroporated though the technique also allows for identifying neurons. We took advantage of the knowledge acquired to show that lentiviral vectors designed with the pCMV promoter infects brain cells and induces egfp expression.

## MATERIAL AND METHODS

### Plasmid DNA preparation

TOP10 E. coli (Invitrogen) were transformed with the plasmid peGFP-C1 (Clontech). Bacteria were grown overnight in lysogeny broth (LB) medium supplemented with kanamycin 1:1000 at 37°C. The plasmid DNA was prepared with the High copy Midi-Prep (Nucleobond Xtra Midi Plus, Macherey-Nagel, Düren, Germany).

### Honeybee electroporation

Worker bees were collected at the hive entrance, were anesthetized on ice, restrained in plastic tubes, fed to satiation with sugar water 30% and left undisturbed. On the next day, they were placed under the binocular and 250 nl of pEGFP-C1 plasmid (Clontech) were injected through the median ocellus following the course of the ocellar tract, directly into the mushroom bodies or directly into the protocerebral lobe. When using the local injection protocols, only small volumes, estimated in the nl range, were injected. Injection was performed with glass capillaries linked to a pneumatic microinjector, by using the method of Müβig and colleagues [9]. Immediately after the injection thin cuts were performed in the cuticle at the dorsal edge of each compound eye and the tip of the electrodes were inserted in the retina of each eye. For the local injection protocols, a window was cut in the cuticle, adjacent tissues were put aside and the electrodes were placed at the surface of the retina. Square pulses of different voltages and various durations were applied at a frequency of 1 Hz (e.g. 25 ms stimulation followed by 975 ms rest). The animals were left to recover after the electroporation and fed with 2 drops of 30 % sugar solution.

### Electrodes

The first experiments were performed with the manufacturer’s electrode model CUY567 (Nepagene, Chiba, Japan; Xceltis, Mannheim, Germany), composed of 2 parallel needles of 0,5 mm diameter. Also, custom made platinum electrodes of different diameters (0.125 mm (large wire electrodes) and 0.05 mm (small wire electrodes), (Advent Research Materials Ltd, Witney, United Kingdom)) with Teflon insulation were used. The wires were soldered on connectors that were plugged into the electroporator. The large wire electrodes were used unmodified (circular section) or as plate electrodes. For the latter, the tips of the electrodes were flattened between 2 metal plates that were hit with a hammer. The Teflon insulation of the wires was removed at the tip by burning it with a lighter. Paper tape was placed to the tip of the wire that was fixed on a holder and moved with a micromanipulator.

### Western blot

Honeybees were anaesthetized on ice and decapitated. Head capsules were fixed on melted wax and brains were dissected. The samples were homogenized in 1X phosphate-buffered saline (PBS), 2 mM Ethylenediaminetetraacetic acid (EDTA), 2 mM ethylene glycol-bis(β-aminoethyl ether)-N,N,N’,N’-tetraacetic acid (EGTA) (PBS-EE) with a Teflon-glass homogenizer. One sample consisted of 2 to 4 brains. Homogenates were sonicated for 10 min and centrifuged at 4°C at 20,000 g for 10 min. The supernatants were discarded and the pellets were re-suspended at a concentration of 1 brain/50 μl in PBS-EE. Sample buffer (0.225 M tris(hydroxymethyl)aminomethane (Tris) pH 6.8, 50% glycerol, 5% sodium dodecyl sulfate, 0.05% Bromophenol blue, 0.25 M dithiothreitol) was added and the samples were sonicated for 10 min and stored at −80°C until analysis. To examine eGFP production, samples were subjected to sodium dodecyl sulfate – polyacrylamide gel electrophoresis (SDS-PAGE) and transferred onto nitrocellulose membranes. The membranes were blocked in 1X PBS, 0,1% Tween-20, 5% non-fat milk powder (blocking solution), 1 hour at room temperature (RT). The membranes were probed with a goat ant-eGFP antibody (1:500, Ab6673, Abcam, Berlin, Germany) dissolved in the blocking solution, overnight at 4°C. The membranes were washed with 1X PBS, 0,1% Tween-20, 4 times for 10 min at RT and then incubated with the secondary antibodies directed against goat IgG coupled to horseradish peroxidase (1:100,000 – Sigma, Munich, Germany) dissolved in the blocking solution, 1 hour at RT. The membranes were washed with 1X PBS, 0,1% Tween-20, 4 times for 10 min at RT and were developed by enhanced chemiluminescence detection (100 mM Tris pH 8.6, 625 μM Luminol, 15 μM p-Coumaric acid, 0.0175 % H_2_O_2_) and the signals were acquired with a LAS1000 camera and the software Image Reader LAS1000 2.60 (Fujifilm, Düsseldorf, Germany).

### Histology and Confocal microscopy

The brains were dissected, dehydrated in an ascending ethanol series and cleared with methylsalicylate, fixed 1.5 hours at RT in 4% paraformaldehyde, 0.2% Triton X-100 in 1x PBS, embedded in 6% low melting agarose and cut into slices of 100 μm with a vibratome Leica VT 1000S (Leica, Bensheim, Germany). The slices were washed twice in 1x PBS, 0.2% Triton X-100 and blocked in 10% normal goat serum (Sigma, Darmstadt, Germany) in 1x PBS, 0.2% Triton X-100. They were stained for 2 days at 4°C with a mouse-derived antibody against GFP (1:400, mAB 3E6, Invitrogen, Darmstadt, Germany), washed for 2 hours at RT in 1x PBS, 0.2% Triton X-100 and incubated overnight at 4°C with a CY5-conjugated secondary antibody (1:400, goat anti-mouse, Jackson Immunoresearch, Suffolk, UK), washed in 1x PBS and embedded in 1x PBS, 80% glycerol and imaged with the Leica TCS-SP2 confocal microscope using a 10x dry objective (HC PL APO CS 10×0.4, Leica, Bensheim, Germany), a 20x dry objective (HC PL APO CS 20.0×0.70 UV), a 40x oil objective (HCX PL APO CS 40.0×1.25) or a 63x oil objective. The Helium Neon (HeNe) 633 nm laser line was used to detect the Cy-5 signal (excitation 650 nm, emission 670 nm) and the Argon/Krypton laser to detect the GFP signal (excitation 488 nm, emission: 507 nm). All brains were imaged at different laser powers lines. Each section was also scanned in bright field, which allowed us to reconstruct the neuropile borders. All pictures were visualised and processed with the software ImageJ (NIH, USA).

### Lentivirus

We tested the functionality of lentivirus developed by the team of Dr. Uwe Maskos (Institut Pasteur, Paris, France) to infect mammalian neurons [21]. The lentivirus expression vector comprises several modifications. It is designed to be replication-incompetent and thus it does not destroy the infected cells. Other modifications enhance infection of non-dividing cells and increase transgene expression of an egfp cassette under the control of the pCMV promoter. Viral particles were generated by co-transfection of HEK-293T cells with the vector plasmid, a packaging plasmid and an envelope plasmid. The viral particles used in our experiments were characterised by a VSV-G envelope protein that allows a broad tropism. Two hundred nl of the viral solutions were injected through the median ocellus directly into the brain of adult honeybees as described above. The animals were sacrificed for histological analysis 3 days post-injection.

## RESULTS AND DISCUSSION

### Specific egfp expression in the brain of electroporated individuals

The efficiency of the electroporation was evaluated with the peGFP-C1 plasmid, expressing egfp under the control of the pCMV promoter. EGFP translation was analysed by Western blot and histology on honeybee brains. A single protein migrating, as expected, at an apparent molecular weight of 32 kDa was detected by Western blot on brains of electroporated animals; but not in animals injected with the plasmidic DNA that were not electroporated (Figure 1A) or animals injected with PBS that were electroporated (data not shown). There was an increase in the amount of eGFP during the first 3 days following the electroporation (Figure1A). EGFP production was confirmed by histological analysis on slices of electroporated brains. The specificity of the signals was demonstrated by showing that the brains of electroporated animals emit native eGFP and Cy-5 signals, the latter being specific of the secondary antibody used for eGFP immunodetection (Figure 1B, C). In addition, scanning the slices at a non-relevant wavelength, specific for Cy3, did not yield any signal (Figure 1B) and the eGFP and Cy-5 signals spatially overlap (Figure 1C). The Cy-5 signals were of greater intensity than the native eGFP signals. For this reason, only the former is presented in the figures. The immunohistological analysis confirmed that eGFP production increased from day 1 to day 3 post-injection (Figure 1D). EGFP signals on day 1 were scarce and signals of low intensity were detected. Most signals were detected in somata regions. After 3 days, the eGFP-positive areas increased and signals of higher intensity were detected mostly in the cortical region of the different neuropiles composing the brain. This shows that cells that were effectively electroporated were scattered within the brain and eGFP production increased during the first 3 days following the electroporation within positively stained cells.

**Figure 1:**
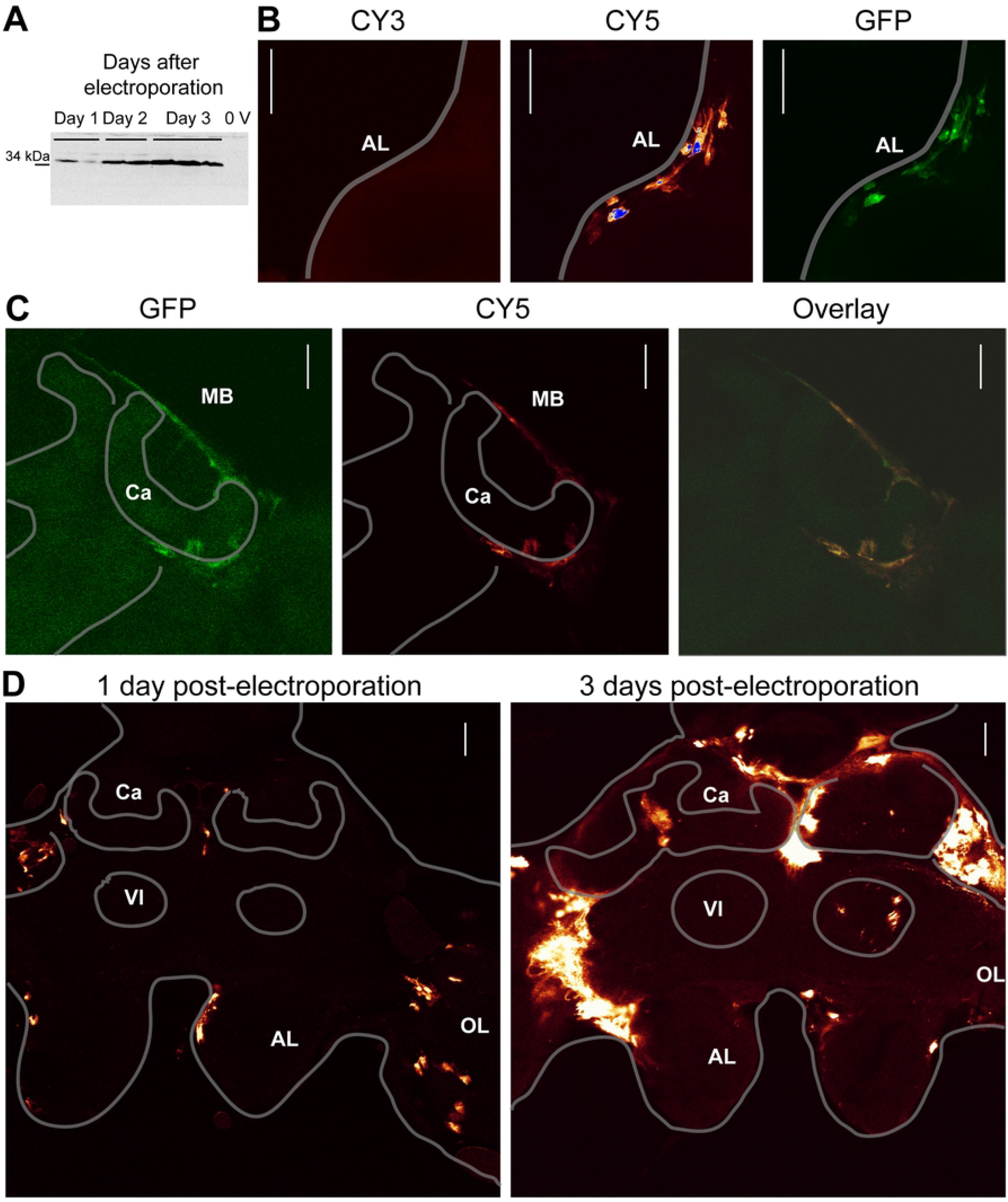
Specific egfp expression in the brain. **A**: Extract from honey bee brains 1 to 3 days after electroporation analysed by Western blot. All bees were injected with pEGFP-plasmid through the median ocellus, electroporation was carried out at 80V with 5×25ms pulses with the large wire electrodes and brains were analysed 1 (Day 1), 2 (Day 2) and 3 days (Day 3) after the electroporation. For the negative control (**0 V**) bees were injected with plasmid, but not electroporated and sacrificed after 1 day. **B, C**: Histological analyses of brain sections scanned at different wavelengths specific for native eGFP signals (GFP) and for the secondary antibody (CY5) or non-specific (CY3). In **B,** the plasmid was injected through the median ocellus and the electroporation was carried out with the large wire electrodes placed on the AL at 10 V with 3×50 ms. The animals were scarified 2 days after the electroporation. In **C**, the plasmid was injected through the median ocellus and the animals were electroporated with the large wire electrodes at 100V with 5×25ms and sacrificed 1 day post-electroporation. The native eGFP and the CY5 signals overlay. **D.** Immunohistological analysis of eGFP signals 1 and 3 days post-electroporation. The plasmid was injected through the median ocellus and the electroporation was carried out with the plate electrodes at 100 V with 5×50ms (3 days post electroporation) and at 80V with 5×25ms (1 day post electroporation). **B, C** and **D**: Section Z-axis 1 μm, scale bar: 100 μm. **AL:** antennal lobe, **MB** : mushroom body **Ca**: calyx, **Vl**: ventral lobe, **OL**: optic lobe.

### Comparison between protocols

To determine the optimal transfection conditions, different electrodes and injection protocols were compared. First, we evaluated the mortality one day after the electroporation with the CUY567 electrode and the large wire electrodes. Potentials ranging from 30 V to 250 V, varying in number and duration of pulses were applied (Table 1). No bees died at potentials of 30 V and 50 V and the mortality rate induced at 70 V was similar to non-electroporated animals. At a potential of 100 V the mortality rate increased to 26 %, it was of 41 and 49 % at 130 V and 150 V, of ~63 % at 170 V, 86 % at 200 V and 100% at 250 V. In some conditions (7 pulses, 25 ms, 100 V, mortality 13%; 3 pulses, 50 ms, 150 V, mortality 7%), *very* low mortality rates were observed compared to other categories at the same potential. These conditions were probably artefacts due to the size of the CUY567 electrode whose positioning was not well controlled. For this reason, custom Teflon insulated electrodes (large wire electrodes) were tested. The mortality induced with these electrodes was higher (33 % at 50 V, 27 % at 70 V, 59 % at 100 V, 64 % at 130 V and 81 % at 140-150 V). However, their positioning could be better controlled, and thus they were preferred to the CUY567 electrode.

**Table 1.** Mortality induced with the CUY567 and the large wire electrodes by varying number of pulses, pulse duration and voltage one day after the electroporation.

| Voltage (V) | Pulses | Duration (ms) | Bees | Mortality (%) |
| --- | --- | --- | --- | --- |
| 0 | 3 | 25 | 24 | 8 |
| 30 | 3 | 25 | 10 | 0 |
| 50 | 3 | 25 | 10 | 0 |
| 70 | 3 | 25 | 18 | 11 |
| 100 | 3 | 25 | 25 | 32 |
| 100 | 3 | 50 | 24 | 33 |
| 100 | 5 | 50 | 29 | 24 |
| 100 | 7 | 25 | 15 | 13 |
| 130 | 3 | 25 | 17 | 41 |
| 150 | 3 | 25 | 15 | 40 |
| 150 | 3 | 50 | 15 | 7 |
| 150 | 5 | 25 | 123 | 58 |
| 150 | 5 | 50 | 30 | 80 |
| 150 | 7 | 15 | 15 | 60 |
| 170 | 5 | 10 | 21 | 62 |
| 170 | 5 | 15 | 23 | 65 |
| 170 | 5 | 25 | 63 | 57 |
| 170 | 7 | 15 | 21 | 68 |
| 200 | 3 | 25 | 10 | 100 |
| 200 | 5 | 25 | 11 | 73 |
| 250 | 3 | 25 | 10 | 100 |

**Large wire electrode**
| Voltage (V) | Pulses | Duration (ms) | Bees | Mortality (%) |
| --- | --- | --- | --- | --- |
| 50 | 5 | 25 | 15 | 33 |
| 70 | 5 | 25 | 15 | 27 |
| 100 | 5 | 25 | 14 | 36 |
| 100 | 7 | 25 | 12 | 67 |
| 100 | 5 | 50 | 15 | 73 |
| 130 | 5 | 25 | 25 | 64 |
| 140 | 5 | 25 | 27 | 82 |
| 150 | 5 | 25 | 35 | 80 |

Increasing the number of pulses or the pulse duration at given potentials did not reveal a marked effect on mortality rate probably because the range of variation of these parameters was too small (Table 1).

The permeabilization of the cell membrane depends on the field intensity around the 2 electrodes. The field strength is proportional to the distance between the electrodes and is expressed in V/cm, but it is also influenced by the diameter and the form of the electrode. Therefore, using different electrodes to generate different electric fields between the electrodes can improve the efficiency of the electroporation [22] [23]. Thus, large and small wire electrodes and the plate electrodes were tested in different protocols by applying 5 pulses of 25 ms and potentials ranging from 80 V to 100 V (Table 2). The applied potentials were adapted to generate calculated field potentials of 200-300 V/cm. The mortality increased during the first 3 days following the electroporation. The animals did not resist direct current injections into the protocerebral lobe. Then the highest effect on mortality was seen after electroporation using the plate electrodes after ocellar injection (71 % 3 days post-injection) and the lowest with the small wire electrodes (30 % 3 days post-injection) (Table 2).

**Table 2:** Mortality rates after electroporation using different protocols.

| Protocol | n | Mortality rate after electroporation (%) |  |  |
| --- | --- | --- | --- | --- |
|  |  | day 1 | day 2 | day 3 |
| Large wire electrodes, ocellar injection | 33 | 15 | 24 | 39 |
| Plate electrodes, ocellar injection | 87 | 35 | 55 | 71 |
| Plate electrodes, mushroom body injection | 52 | 31 | 46 | 54 |
| Small wire electrodes, ocellar injection | 40 | 13 | 25 | 30 |
| Large wire electrodes, protocerebral lobe injection | 7 | 100 | - | - |

All brains were electroporated by applying potentials ranging from 200 to 300 V/cm.

The efficiency of the electroporation was evaluated in transfected animals, either by Western blot (data not shown) or by immunohistology. The transfection efficiency ranged from 70 % to 92 % of the electroporated brains (Table 3). The electroporation did not result in homogeneous transfections of all cells but varied between individuals and protocols. Ocellar injections resulted almost always in egfp expression in the ocellar tract. The mushroom body (MB), the antennal lobes (AL) and the optic lobes (OL) were also frequently transfected, whereas cells of the protocerebral lobe (PL) were rarely transfected. Injection directly into the MB led to egfp expression mostly but not exclusively in this neuropile.

**Table 3:** EGFP distribution 3 days after electroporation (200-300 V/cm).

| Protocol | n | GFP (%) | GFP-positive regions (%) |  |  |  |  |
| --- | --- | --- | --- | --- | --- | --- | --- |
|  |  |  | AL | OL | MB | OT | PL |
| Large wire electrodes, ocellar injection | 13 | 92 | 67 | 67 | 75 | 100 | 17 |
| Plate electrodes, ocellar injection | 29 | 76 | 59 | 45 | 73 | 100 | 27 |
| Plate electrodes, mushroom body injection | 20 | 70 | 14 | 29 | 93 | 14 | 14 |
| Small wire electrodes, ocellar injection | 19 | 79 | 67 | 53 | 67 | 93 | 27 |

AL: antennal lobes; OL: optic lobes; MB: mushroom bodies; OT: ocellar tract; PL: protocerebral lobe. The percentages of GFP-regions are calculated relatively to the total amount of GFP-positive brains.

EGFP signals induced with the different protocols were disparate and localised most of the time in cortical areas (Figure 2A, 2B). In some brains the calycal neuropile was also stained (Figure 2B, 2C). Although it was also detected in several neuropiles such as the AL, and the PL, the applied protocols resulted in expression mainly in the MB and the OL regions, probably because the DNA concentration was higher at the injection site and because these neuropiles were closer to the electrodes. The large wire electrodes combined with ocellar injections induced the highest rate of transfections in the different neuropiles (Table 3, Figure 2A). However, larger areas were transfected with the plate electrodes, which were representing signals of higher intensity (Figure 2B). After local injection into the MB and electroporation with the plate electrodes, the majority of transfected cells were found within this neuropile but were not restricted to it (Figure 2C). The electroporation with the small wire electrode combined with ocellar injections yielded similar results (Figure 2D).

**Figure 2:**
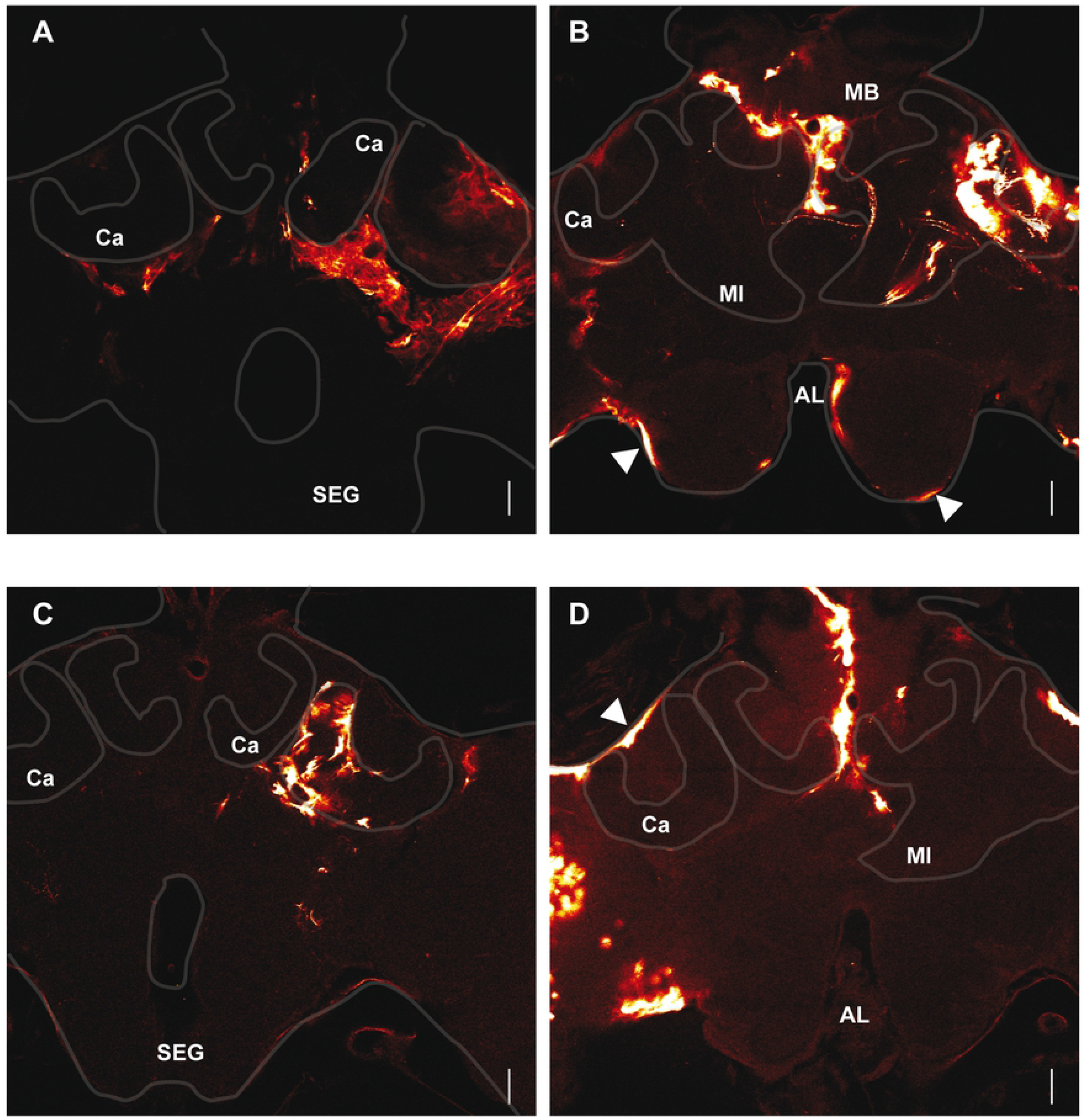
Distribution of anti-eGFP signals 3 days after different electroporation protocols. **A:** The animals were injected through the median ocellus and electroporated with the large wire electrodes. 100V, 5x 25 ms. **B:** The animals were injected through the median ocellus and electroporated with the plate electrodes. The white arrowheads indicate perineural surface glia cells. 100 V, 5x 50 ms. **C:** The animals were injected into the MB calyx and electroporated with the plate electrodes. 100 V, 5x 25 ms. **D:** The animals were injected through the ocellus and electroporated with the small wire electrodes. The white arrowheads indicate subperineural surface glia cells. 100 V, 5x 25 ms. **AL:** antennal lobe, **SEG**: suboesophageal ganglion, **Ca**: calyx, **Ml**: medial lobe. Section Z-axis 1 μM, Scale bar: 100 μm.

### Transfected cell types

It was not possible to determine the morphology of single cells in most of the electroporated brains because cell clusters were transfected, and single cell morphology could not be differentiated. In some cases, clusters composed of a limited number of cells were transfected and it was possible to describe their cellular morphology. The electroporation resulted in the transfection of both neuronal and non-neuronal cells; the latter being transfected more frequently. The observation of the signals shows that the cells are completely stained. Based on their localisation and their morphology, the non-neuronal cells correspond to glia cells, although no cell specific glia markers were tested. Several glia cells types were observed as previously reported in the honeybee [24]. They are located at the surface of the brain, associated with somata or with neuropile regions. The nomenclature for glial cells used by Awasaki and colleagues (2008) for the Drosophila brain will be applied in the following.

Surface glial cells within the brain neurolemma were transfected (Figure 2B, 2D). First, large size cells located at the surface of the brain might correspond to subperineural glia cells (Figure 2D). Smaller slender oblong cells were also observed and might correspond to perineurial glia cells (Figure 2B). Together, the perineural and the subperineural glia cells form the neurolemma [25].

In the cortical layers of the MB, cortex glia cells characterised by a mesh-like morphology that enwrap individual Kenyon cells (KC) somata outside and inside the calyces expressed egfp (Figure 3A). Although individual glial cells could not be distinguished, it is evident that these cells enwrap different KCs subpopulations to build a mesh. In some preparations eGFP positive cell bodies corresponding to KC soma are identified and are integrated in the mesh of cortex glia (Figure 3A). Interestingly, a thick cellular structure built a sheet around the external KC soma layer (Figure 3A). This cellular structure segregated laterally external KC of the collar and the lip from the soma of adjacent neuropiles and extended to the dorsal part of the MB. It seems to emanate from cortex glia cells, although, it cannot be excluded that it is built by surface glia cells closely associated with cortex glia cells. Cortex glia cells were already described in honeybee drone [24], in Drosophila [26] [25] and in the cricket *Acheta domestica* [27] where it was shown that they enwrap individual KC somata. We also observed this cell type in other neuropiles like in the dorsal soma rind of the medulla in the OL or in the median soma rind of the AL (Figure 3Bi, 3Eii). Similar glial cells were described in *Manduca sexta* [28], in Drosophila [25], [29] and in *Musca domestica* [30], for review see Edwards and Meinertzhagen [31].

**Figure 3:**
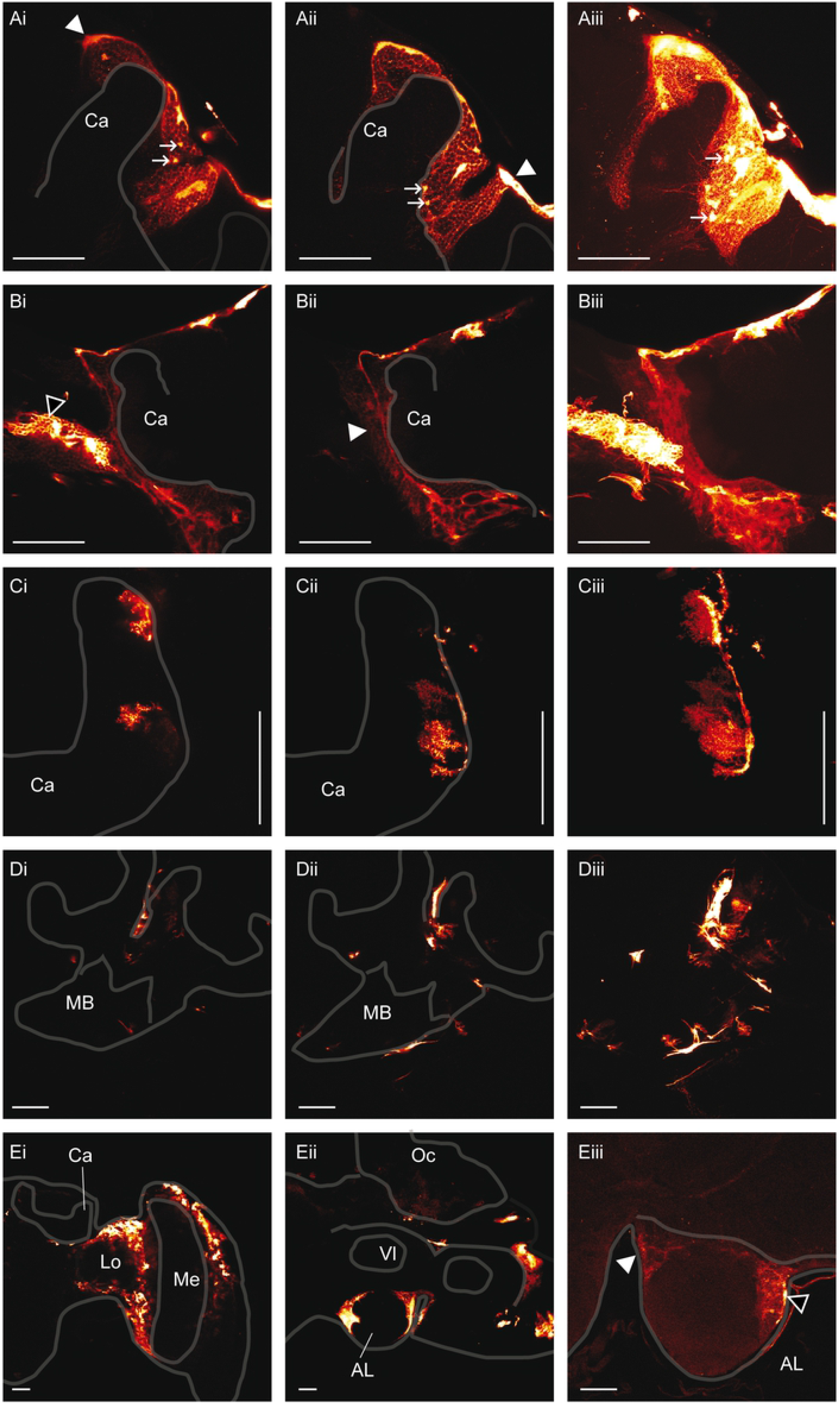
Non-neuronal transfected cell types: **A: Anti-eGFP signals in cortex glia cells within the calycal region of the MB**. The white arrowheads indicate the thicker cellular structure observed at the surface of the brain and probably originating from subperineural glia cells. The white arrows indicate eGFP positive cell soma. The brain was electroporated 5x 25 ms at 80 V with the small wire electrodes and the plasmid was injected in the ocellar tract. The brain was prepared 3 days post-electroporation. **i, ii**: Section Z axis 0.5 μm, **iii:** Assembly of confocal sections along the z-axis over 98 μm depth. **Ca:** calyx. Scale bar: 100 μm **B: Anti-eGFP signal in cortex glia cells of the outer calycal region:** The cell (white arrow head) is observed on the same preparation as a cortex glia of the optic lobe (black arrow head). The brain was electroporated with the large wire electrode 5x 25 ms at 100 V and the plasmid was injected in the ocellar tract **i, ii**: Section Z axis 1 μm, **iii:** Assembly of confocal sections along the z-axis over 46 μm depth. **Ca:** calyx. Scale bar: 100 μm. **C. Anti-eGFP signal in dendritic glia cells of the calyx:** The brain was electroporated with the plate electrode 5x 25 ms at 100 V. **i, ii:** Section Z axis 0,5 μm, **iii**: Assembly of confocal sections along the z-axis over 13 μm depth. **Ca:** calyx. Scale bar: 100 μm. **D. Anti-eGFP signal in fibrous lamellar glia cells of the MB:** The plasmid was injected directly into the MB and **t**he brain was electroporated with plate electrodes 5x 25ms at 100 V. **i, ii:** Section Z axis 1 μM. **iii:** Assembly of confocal sections along the z-axis over 89 μm. Scale bar: 100 μm. **E. Anti-eGFP signal in cortex glia cells of the OL (i) and the AL (ii, iii): i.** The brain was electroporated with the plate electrodes 5x 25 ms at 100 V. Section Z axis 1 μm. **ii.** The brain was electroporated with the plate electrodes 5x 25 ms at 100 V and the animal was injected through the median ocellus. Section Z axis 1 μm. **iii.** The brain was electroporated with the small wire electrodes 5x 25 ms at 100 V. The white arrowhead indicates a cortex glia cell and the black arrowhead an astrocyte-like glia cell. **Ca:** calyx, **Lo:** lobula, **Me:** medula, **Oc:** ocellus, **Vl:** vertical lobe. Section Z axis 1 μm. Scale bar: 100 μm.

A single cortex glia cell in the MB outer cortex was found that exhibits a different morphology (Figure 3B). It is characterised by a long slender fibre that penetrates the cortex between the MB calyx and the OL and extends deeply into the somata layers down to the region delimited by the basis of the calyx, the optic lobe and the protocerebral lobe where it becomes thicker probably to form the cell body. Several processes extend ventrally from the cell body and build a lose network in the cortical area that enwraps cell bodies or groups of cell bodies (Figure 3Bi, 3Bii, 3Biii). Primary and secondary ramifications emanate along the whole central arbor, they extend through the cortical area and also enwrap neuronal cell bodies. The delimitation of the cell bodies is less pronounced as for the cortex glia cells of the inner calycal cortex. The processes extend through MB somata and maybe also through the somata of adjacent neuropile. It would be interesting to determine if the central arbor of the glia cell segregate MB somata from the somata of adjacent neuropiles. A thicker cellular structure is observed at the border of the calycal neuropile but not at the border of adjacent neuropiles (Figure 3B). Overall, these cortex glia cells of the outer calycal layer build a fibrous mesh structure that occupies at least the whole outer MB cortex. The central arbor prolongs dorsally to the surface of the brain, where it bifurcates and extend at the surface of the MB to envelop it. However, it cannot be excluded that the cortex glia cell connects surface glia cells that builds the sheet at the surface of the brain.

Neuropile glia cells were also transfected. These cells have their cell body on the external side of the MB calyx and extend dendritic processes within the calycal neuropile (Figure 3C). They develop from an elongated structure extending to the surface of the neuropile. This cell type was already described in *Drosophila* and in the honeybee as astrocyte-like glia [24, 25]. The cell bodies are located at the outer surface of the neuropile and a dense arborisation penetrates into the neuropile. It is associated with synaptic regions between neurones. This cell type was also observed in the AL though less frequently, probably because our method electroporated predominantly cell somata of the superior protocerebrum (Figure 3Eiii). The other type of neuropile glia cells, the fibrous lamellar type, was also electroporated. It is associated to the surface of the neuropile and delimits its different sub-compartments. It was observed in the MB calyx and in the peduncle (Figure 3D). This cell type was already described in the honeybee [24] and in *Drosophila* [25].

Neuronal cells were efficiently electroporated but only occasionally. One reason for the lack of neural transfections might be that their somata are relatively small and that the electroporation conditions were not optimal to efficiently transfect them. Non-neuronal cell somata are bigger and are therefore probably more susceptible to electroporation. Unfortunately, the electroporation parameters used could not be increased further (max 100 V) due to a too high mortality. Interestingly, in some cases the signals emitted by glia cells of the MB were of high intensity and might have masked neuron specific signals.

In a few cases several KC were successfully electroporated and eGFP signals were observed in the whole cell (Figure 2B, Figure 4). The signals are predominantly observed, but not restricted, in the MB of the left hemisphere, suggesting that the electroporation is polarised to the cathode side, on the left hemisphere. The lateral calyx shows intensive eGFP signals emitted by glia cells but also by KC because the signals are not confined to the calycal neuropile. Indeed, cell soma, dendrites and axonal structures emit eGFP signals. KC axons projecting to the neck of the peduncle by forming 3 main bundles (Figure 4B, 4C) could be traced down to the terminals in the medial (Figure 4B), and the vertical lobes (Figure 4D). Discrete eGFP positive regions corresponding to KC terminals were detected in both lobes. Their localisations correspond to layers representing the lip, the collar and the gamma lobe [32]. The most ventral signals in the vertical and gamma lobes region correspond to KC class II. The KC axon terminals localised on the medial side of the lobe are connected to the medial calyx and those on the lateral side to the lateral calyx. Indeed, eGFP positive regions are detected within both calyces and are in the proximity of eGFP positive axons projecting to the peduncle. Neurons projecting from the OL were also electroporated. Their axons are located at the lateral MB beneath the calyces (Figure 4B, C, D). Some of them probably project to the contralateral hemisphere (Figure 4B) as already described [33]. Some presynaptic terminals could be observed in the collar region of the medial calyx (Figure 4A), as indicated by the presence of boutons arranged along the thin terminals.

**Figure 4:**
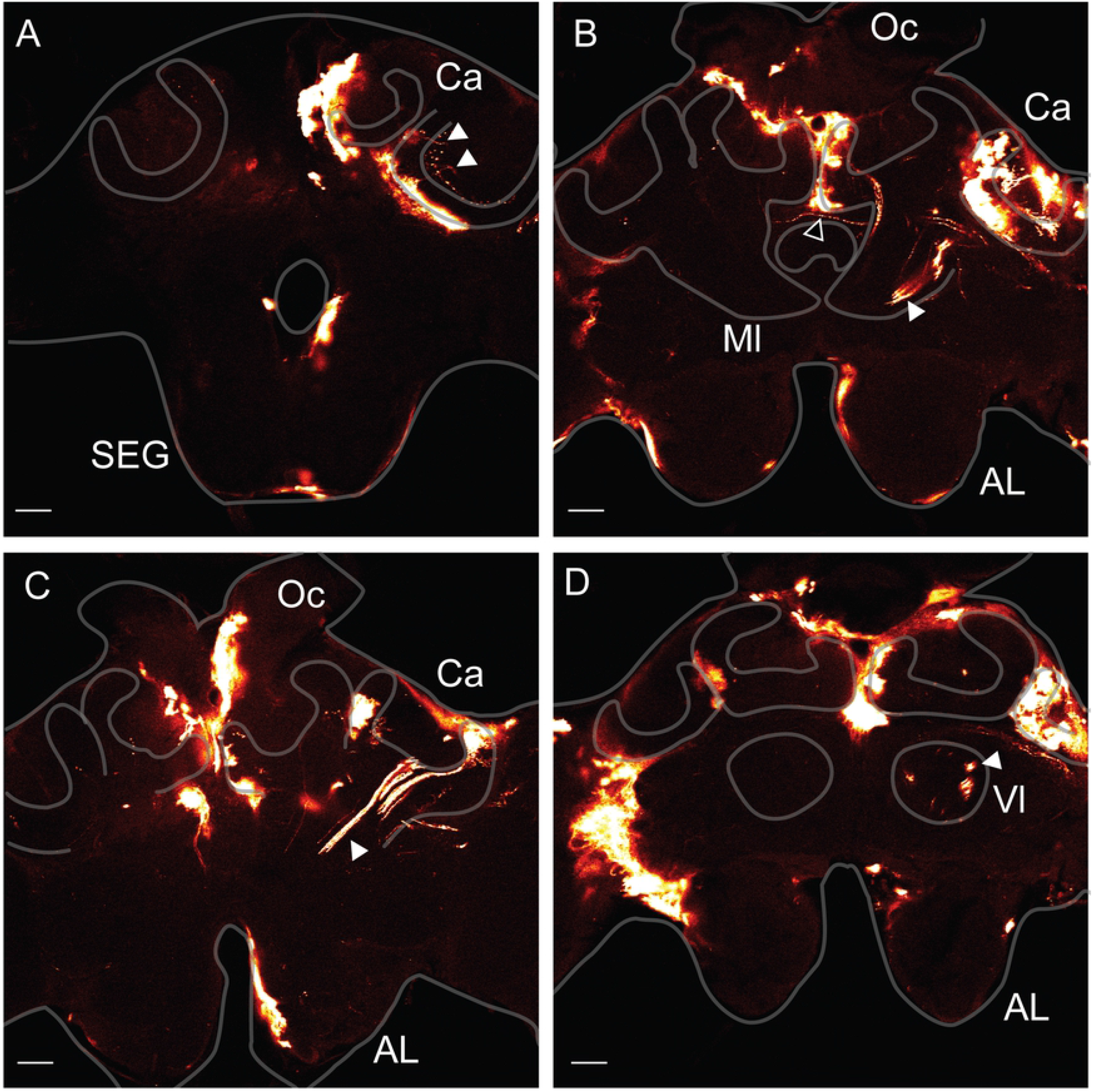
Anti-eGFP signal detected in Kenyon cells: The animals were injected through the median ocellus and electroporated with the plate electrodes at 100 V, 5x 50 ms. **A.** Section of the brain at the level of the suboesophageal ganglion (**SEG**). The white arrow head shows KC somata connected to neurites projecting into the calyx (**Ca**) and to their dendrites. **B.** Section of the brain at the level of the central complex. The white arrow head shows the axons of KC terminating into the median lobe (**Ml**). eGFP positive signals are also detected into the calyx (**Ca**). The black arrowhead shows the axon a neuron crossing laterally the brain. **C.** Section of the brain at the level of the antennal lobe (**AL**). The white arrow head shows KC axons projecting into the neck and the peduncle of the MB. **D.** Section of the brain at the level of the antennal lobe (**AL**) and the vertical lobe (**Vl**). The white arrow head shows KC terminals in the Vl. Scale bar: 100 μm.

The lateral calyx of this brain was visualised at higher magnification (Figure 5A). The cell bodies of several KC class I innervating the lip are localised at the surface of the cortical layer around the neuropile of the lip. Thin dendritic fibres radiate within the neuropile to build broad dendritic domains with thickenings along the fibres probably corresponding to synaptic sites. Figure 5Ai shows the dendritic domains originating from 2 or 3 KC and their axons projecting to the peduncle. In another focal plane, a bigger group of KCs were electroporated. The dendritic domains occupy a large area of the lip and collar neuropile and single dendrites can hardly be differentiated. Accordingly, their axons form a thick bundle projecting to the peduncle (Figure 5Aii, iii). Interestingly, a single KC with its cell body localised within the inner part of the calyx at the surface of the MB was successfully electroporated (Figure 5Aiii). The neurite descending to the peduncle bifurcates first to the collar neuropile. A collar neuron arborizes widely in the same region with thickening distributed along neurites. It is not known if the collar neuron and the KC make synapses. Then the KC neurite exits the collar and descends further to the neck of the peduncle where it enters again into the neuropile at the level of the basal ring to build a discrete dendritic domain (Figure 5Aiii). The proximal part of the axon connecting the cell body cannot be distinguished on figure 5. However, the exposition of the slide to higher excitation showed it (data not shown). Neurons probably originating from the OL also expressed eGFP positive signals. Some of them extend through the whole preparation and cross the MB at the level of the ventral part of the calyx. Two neurites bifurcate from these axons to enter the calycal neuropile and build potentially presynaptic terminals in the collar as characterised by their boutons (Figure 5Ai, ii, iii).

**Figure 5:**
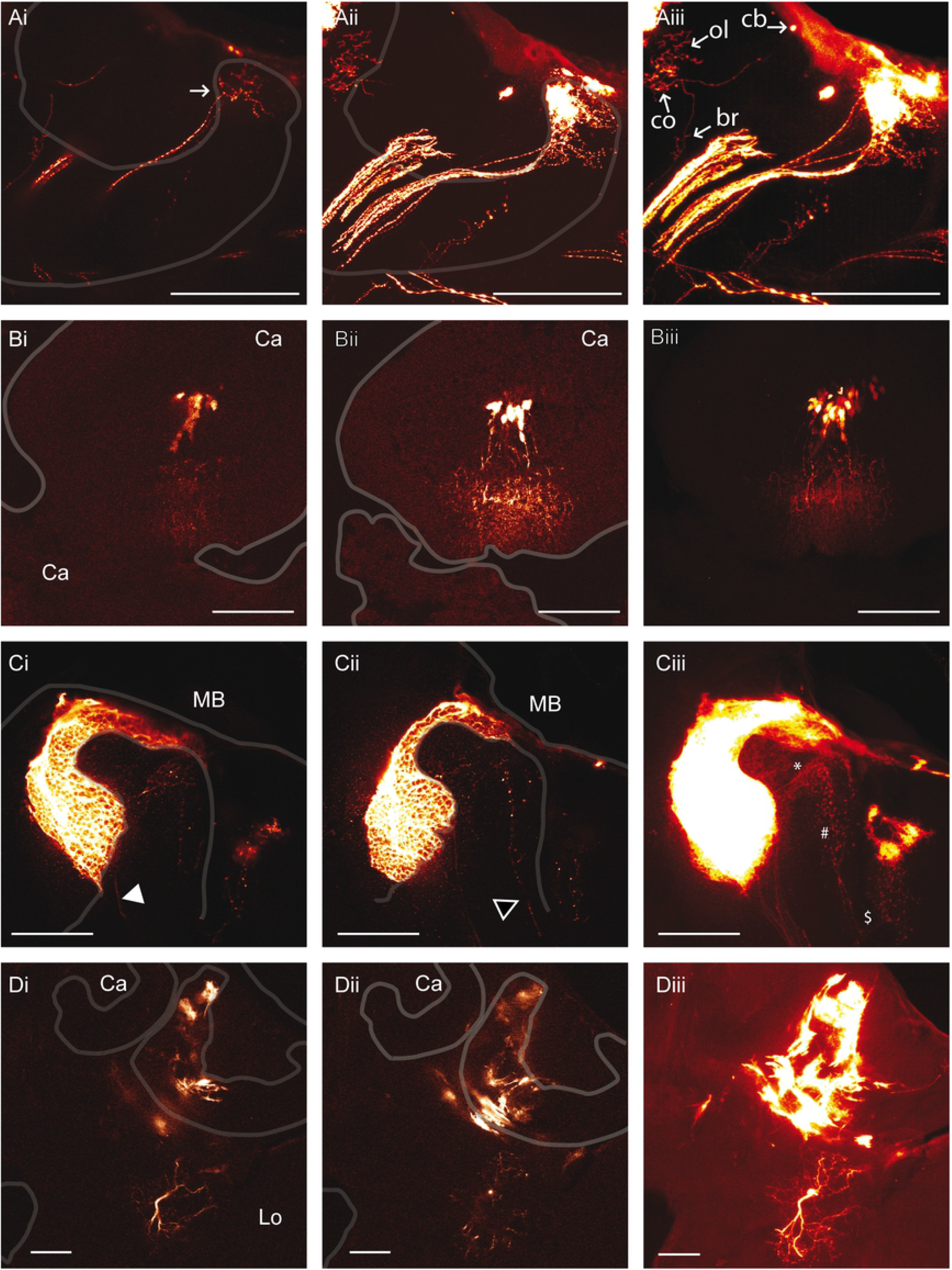
Anti eGFP signals detected in neuronal cell types: **A.** The plasmid was injected through the median ocellus. The brain was electroporated with the plate electrode 5x 25 ms at 100 V. Assembly of confocal sections (0,5 μm) along the z-axis over 31 μm (**i**), over 41 μm (**ii**) and over 135 μm (**iii**). **i.** The white arrow shows a discrete group of KC of the lip. **iii.** The white arrows show the cell body (**cb**), the dendrites of the collar (**co**) and of the basal ring (**br**) of a unique KC. A collar neuron is also stained (ol). **B**. The plasmid was injected through the median ocellus. The brain was electroporated with the plate electrode 5x 25 ms at 100 V. **i, ii**: Section z-axis 1 μm. **iii.** Assembly of confocal sections (1μm) along the z-axis over 85 μm. **C.** The plasmid was injected directly into the MB and the brain was electroporated 5x 25 ms at 100 V with the plate electrodes. The white arrow head shows the axons of KC and the black arrow head the axon of OL neurons entering the MB. **i, ii:** Section z-axis 1 μm. **iii:** Assembly of confocal section along the z-axis over 100 μm. The KC dendrites (**\***), the OL boutons (**#**) and the bifurcation of the axon of the ascending OL neuron ($) are shown. Scale bar: 100 μm. **D.** The brain was electroporated with the plate electrodes 5x 25 ms at 100 V and the plasmid was injected directly into the MB. **i, ii** Section z-axis 1 μm. **iii.** Assembly of confocal sections (1 μm) along the z-axis over 99 μm. **Ca:** calyx, **Lo:** lobula Scale bar 100 μm.

In another preparation, we could also electroporate a group of KCs with its somata located above the neuropile of the basal ring. Thin dendritic structures spread within a big part of the basal ring (Figure 5Bii, iii), the axons form posteriorly a bundle that projects to the peduncle, and then extend to the peduncle (Figure 5Bi, iii).

KCs of the collar characterised by one primary neurite extend almost through the whole neuropile from the inner part to the outer part of the calyx (Figure 5C). They are characterised by thin dendritic processes extending from the central neurite to form a pyramidal shaped dendritic domain becoming broader toward the outer part of the calyx. The axons project to the peduncle. The cell bodies of these KC could not be localised; they probably lie within the inner part of soma layer of the calyx where an eGFP positive glial cell mask them. Peripheral neurons originating from the OL were also observed with their axons that penetrate into the calyx. They harbour thicker bouton-type structures at their terminals corresponding to presynaptic structure participating in forming microglomeruli within the neuropile. Interestingly, before the OL axons enter the collar, axons bifurcate, one part enter the medulla of the OL and form broad ramifications that also exhibit thicker structures corresponding to presynaptic terminals. The cell body of these cells could not be identified.

Figure 5D shows a neuron of the protocerebral lobe that is located beneath the lateral calyx. It is composed of a primary thick neurite (Figure 5Di, iii) that emanates from the cell body. It probably corresponds to the axon extending ventrally and bifurcating in 2 branches, which harbour thinner ramifications (Figure 5Di, iii). Several thin neurites, probably corresponding to dendrites, extend dorsally from the cell body and come close to the calycal neuropile without penetrating it (Figure 5Di, iii).

### Egfp expression via lentivirus transduction

The knowledge acquired during the electroporation experiments allowed the design of lentiviruses expressing egfp under the control of pCMV. Lentivirus-based expression systems provide several advantages over other virus-based expression strategies. They are retroviruses capable of stable integration into the genome of the host cell and they are capable of genomic integration into non-dividing cells, such as neurons. We tested in the honeybee the functionality of lentivirus characterised by a VSV-G envelope protein that allows a broad tropism. It was shown that they can transduce neurons in mice [21]. Here, the viral solutions were injected through the median ocellus directly into the brain of adult honeybees. There was no noticeable mortality induced by the injection of viruses 5 days after the injection. EGFP was observed in many cell bodies surrounding the calycal and the suboesophageal ganglion neuropiles (Figure 6). The transduced cells are most probably glia cells associated with the shaping of the neuropile because staining is found only in regions surrounding the neuropiles and not in neuropiles. Although no expression was observed in neurons, this result proves the functionality of the system in the honeybee. It opens the gate to the design of other viral vectors, for example characterised by another envelope, that will result in the transduction of neurons.

**Figure 6:**
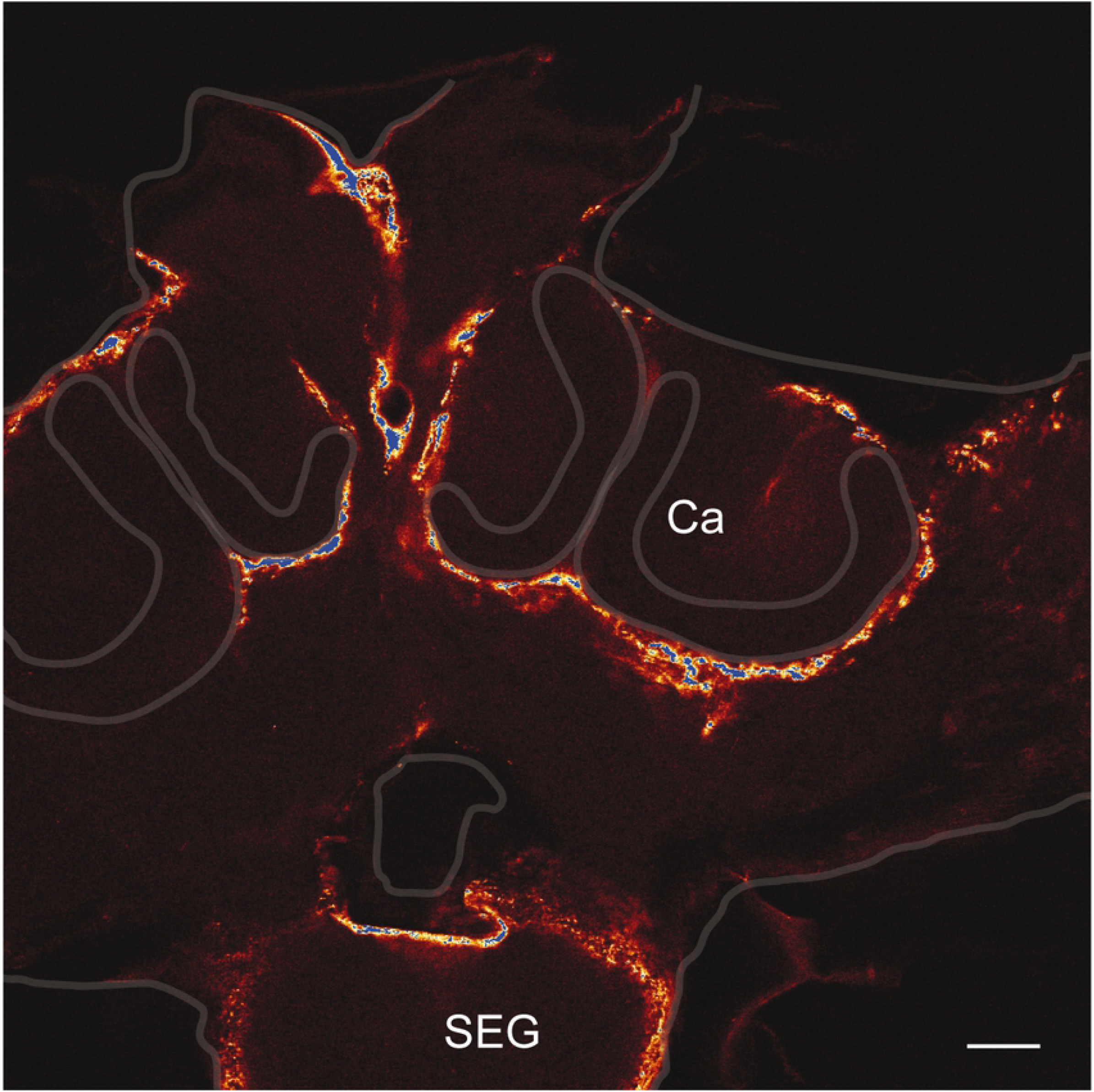
Anti-eGFP signals in bees injected with Lentiviruses. The bees were injected with viral particles through the median ocellus and the brain was dissected 3 days post-injection. Calyx (**Ca**), Suboesophageal ganglion (**SEG**). Scale bar 100 μm.

## CONCLUSION

In this study we re-evaluated the potential of in-vivo electroporation to induce recombinant protein production with the peGFP-C1 plasmid in the honeybee brain. Different protocols were applied and led principally to egfp expression in glia cells but also in neurons of different types. The technique was designed to electroporate cells principally in the dorsal part of the brain and indeed, most electroporated cells were localised in the MB and the dorsal part of the OL, although not restricted to these neuropiles. The electroporation did not result in the transformation of homogeneous cell populations. Thus, the technique is so far not adapted to manipulate brain physiology for behavioural studies. However, it is potentially suitable for anatomical and optical imaging studies, for example with the calcium sensor Gcamp [34]. We describe cortex glial cell types in the outer cortical layer of the calyx that were to our knowledge not yet described in insects, based on their particular morphology.

The viral expression system allowed a more homogenous transduction of cell population. However, it was expected to target neurons as in mammals, but instead glia cells were transduced. Although transgenic animals and the CRISPR/Cas9 techniques constitute the actual state of the art, the techniques evaluated in this study remain of high importance, because they are more adapted for the manipulation of somatic cells only, and are not inherited, which is safer for behavioural and physiological studies because it allows the release of animals into the environment.

## ACKNOWLEDGMENTS

This work was supported by the Deutsche Forschungsgemeinschaft (DFG) through grant no. LE1809/1-1 and LE1809/1-2. We thank Dr. Jürgen Rybak for the critical review of the manuscript.

## ABBREVIATIONS

egfp (gene) / eGFP (protein): enhanced green protein
pCMV: cytomegalo virus promoter
PBS: Phosphate-buffered saline
PBS-EE: 1X PBS, 2 mM Ethylenediaminetetraacetic acid
MB: Mushroom body
AL: antennal lobes
OL: optic lobes
PL: protocerebral lobe
KC: Kenyon cell

